# Effects of CwlM, a peptidoglycan synthesis regulator, on beta-lactam resistance and host-pathogen interactions

**DOI:** 10.1101/2025.07.08.663695

**Authors:** Cátia Silveiro, Diana Mortinho, Francisco Olivença, Manoj Mandal, David Pires, Elsa Anes, Maria João Catalão

## Abstract

**Background:** The emergence and spread of multidrug-resistant (MDR) strains of *Mycobacterium tuberculosis* (*Mtb*) urge the development of novel drugs and efficient therapeutic programs. A recent study aiming to uncover differential beta-lactam susceptibility phenotypes in clinical strains of *Mtb* found that the M237V substitution in *cwlM* (*Rv3915*) was associated with increased susceptibility to amoxicillin. Considering that *Mycobacterium smegmatis* (*Msm*) is a widely used surrogate model for *Mtb*, we constructed a *cwlM* knockdown mutant in *Msm* using CRISPR interference (CRISPRi) to elucidate the role of CwlM in beta-lactam susceptibility and intracellular survival.

**Results:** Quantitative RT-PCR assays confirmed the successful repression of *cwlM*, while the phenotyping assays confirmed the essentiality of CwlM-related processes for mycobacterial viability. Collectively, the antibiotic susceptibility assays suggested that CwlM_SMEG_ may promote beta-lactam resistance, particularly to meropenem and cefotaxime. Moreover, CwlM_SMEG_ was found to support *M. smegmatis* intracellular survival within THP-1-derived macrophages. To address conflicting reports regarding its predicted peptidoglycan (PG) hydrolase activity, we purified recombinant CwlM_TB_. The *Micrococcus luteus*-derived PG-based zymogram indicated that CwlM_TB_ lacks PG-hydrolytic activity, suggesting it might act as a regulator of PG biosynthesis instead.

**Conclusions:** Our findings indicate that CwlM contributes to beta-lactam resistance and intracellular survival, regardless of lacking detectable PG-hydrolytic activity. Overall, CwlM was found to be essential and highly vulnerable, highlighting its potential as a therapeutic target that warrants further investigation.

## Background

Tuberculosis (TB) remains the deadliest infectious disease worldwide, having claimed 1.25 million lives in 2023 and causing over 10 million TB cases annually, a rising trend since 2021 [1]. The emergence and dissemination of multi- and extensively-drug-resistant (MDR/XDR) TB urge the identification of novel drug targets for the development of effective curative treatments [1,2]. In this context, a structure worth exploring is the peculiar cell wall (CW) of mycobacteria, which is essential for its viability and virulence. The mycobacterial CW encompasses a distinctively cross-linked and enzymatically modified peptidoglycan (PG), a highly branched arabinogalactan (AG) polysaccharide and characteristic long-chain mycolic acids (MA) [3,4].

Despite its uniqueness, the mycobacterial PG is still overlooked when identifying antitubercular targets. The biosynthesis of PG starts in the cytoplasm with the generation of uridine diphosphate (UDP)-*N*-acetylglucosamine, which is then converted into UDP-*N*-acetylmuramic acid (UDP-Mur*N*Ac) by MurA and MurB [3,4]. After, the *N*-acetylmuramic acid hydroxylase NamH catalyzes the hydroxylation of UDP-Mur*N*Ac to UDP-*N*-glycolylmuramic acid (UDP-Mur*N*Glyc) [5]. This PG modification promotes resistance to lysozyme and beta-lactams [5,6] and modulates host-pathogen interactions [6-9]. The subsequent addition of L-Ala (MurC), D-Glu (MurD), *meso*-DAP (MurE) and D-Ala-D-Ala (MurF) produces the muramyl-pentapeptide, which is then anchored to the plasma membrane (PM) by MraY/MurX, generating lipid I [3,4]. The final intracellular step of PG synthesis, the formation of lipid II, is catalyzed by MurG. Both lipid I and II are then amidated at the α-carboxyl group of D-*iso*glutamate (D-*i*Glu) by the MurT/GatD amidotransferase complex [10-12]. The amidation of D-*i*Glu is essential for PG cross-linking in mycobacteria [13,14], modulates PknB-dependent cell division [15], promotes resistance to beta-lactams and lysozyme [6,14], and functions as an immune evasion strategy during infection [6,9]. Likewise, the amidation of lipid II at the ε-carboxyl group of *m*-DAP is catalyzed by AsnB [16,17]. Apart from being essential for cell growth and L, D-transpeptidase (Ldt) activity in *Mycobacterium tuberculosis* (*Mtb*) [18], the amidation of *m*-DAP also modulates resistance to antibiotics and lysozyme [19,20] as well as host immune responses. Once the MurJ flippase catalyzes the translocation of modified lipid II across the PM, the PG is polymerized by transglycosylases, penicillin-biding proteins (PBPs) and Ldts.

Beta-lactams, a broadly employed and reliable antibiotic class, could be incorporated into alternative drug-resistant TB (DR-TB) therapeutic schemes, offering improvements over the current lengthy and toxic regimens. These regimens usually exclude beta-lactams due to the intrinsic resistance of *Mtb* to these antibiotics, credited not only to a potent beta-lactamase [21] but also to the impervious layer of MA and the predominance of non-classical Ldt-catalyzed 3→3 cross-links [22,23]. Nonetheless, several reports found that MDR-TB strains are particularly susceptible to carbapenems when combined with beta-lactamase inhibitors, such as clavulanate [24-29]. A previous study screened 172 clinical strains of *Mtb* for differential beta-lactam susceptibility phenotypes and conducted a whole genome sequencing analysis to identify cases in which the prospective addition of beta-lactams to TB therapy may yield added benefits [29]. The M237V substitution in *cwlM* (*Rv3915*), present in 50/172 isolates, was associated with increased susceptibility to amoxicillin, with and without clavulanate [29].

Being highly conserved amid mycobacteria and essential for *Mtb* survival, CwlM is predicted to be a *N*-acetylmuramoyl-L-alanine amidase, a type of PG hydrolase [30,31]. Whereas Deng et al. [30] have shown that CwlM_TB_ possesses autolysin activity, a recent study found the function of CwlM to be regulatory rather than enzymatic [31]. CwlM is the main substrate of the essential serine/threonine protein kinase (STPK) PknB [32]. Replicating *Mtb* produces two forms of CwlM: phosphorylated CwlM binds to FhaA, a fork head-associated domain protein, whereas non-phosphorylated membrane-associated CwlM interacts with the MurJ flippase [32]. In nutrient-replete conditions, CwlM is phosphorylated in the cytoplasm, binds to MurA, the first enzyme in PG biosynthesis, and stimulates its catalytic activity by _∼_30-fold [31]. In starvation or stasis, CwlM is dephosphorylated (possibly by PstP) [33], consequently downregulating PG biosynthesis and promoting antibiotic tolerance [31]. In this case, the activity of MurJ is stimulated, resulting in the accumulation of non-incorporated PG muropeptides in the periplasm [32]. Overall, CwlM simultaneously regulates the biosynthesis of PG precursors and their translocation across the PM. Therefore, the balance between both forms of CwlM, influenced by altered PknB expression or activity, is essential for cell viability [32].

Here, we implemented the dCas9_Sth1_-based CRISPRi system [34] in *Mycobacterium smegmatis*, a widely used surrogate for *Mtb*, to uncover the role of the PG hydrolase homolog, CwlM_SMEG_, in susceptibility to beta-lactams and host-pathogen interactions. Additionally, we purified recombinant CwlM_TB_ to elucidate its activity.

## Methods

### Bacterial strains, culture conditions, plasmids, cell lines, and antibiotics

A description of the bacterial strains, plasmids, and cell lines employed in this study is provided in **Table 1**. *Escherichia coli* (*E. coli*) strains were grown in Luria-Bertani (LB) media (Merck), overnight (ON) at 37°C, to amplify the CRISPRi backbone PLJR962 (Addgene #115162) and for cloning. *Micrococcus luteus* (*M. luteus*) strains were also grown in Luria-Bertani (LB) media and used for the zymography assay. *M. smegmatis* strains were grown in Middlebrook 7H9 broth (BD™ Biosciences) supplemented with 0.2% glycerol (ThermoScientific), 0.05% tyloxapol (Sigma-Aldrich), and 0.5% glucose (Sigma-Aldrich). For plating procedures, mycobacteria were grown on 7H10 agar (BD™ Biosciences) with 0.5% glycerol. Bacteria were incubated at 37°C with (liquid cultures) or devoid of (plates) shaking at 180 rpm. Media for all bacteria were supplemented with 25 μg/mL of kanamycin as appropriate. To stimulate target gene knockdown by the dCas9_Sth1_-sgRNA complex, 100 ng*/*mL of ATc were added to the cultures when necessary. RPMI-1640 media (Gibco) supplemented with 10% fetal bovine serum, 1% sodium pyruvate, 1% L-Glutamine, 1% HEPES buffer and 1% non-essential amino acids was used to grow THP-1 cells at 37°C, with an atmosphere of 5% of CO_2_. Antibiotic stocks of amoxicillin (AMX), cefotaxime (CTX), ethambutol (EMB), isoniazid (INH), meropenem (MEM) and vancomycin (VAN) (all from Sigma-Aldrich) were prepared in purified MilliQ water to a final concentration of 1.28 mg/mL. The stocks of ATc (Sigma-Aldrich) were prepared in dimethyl sulfoxide (DMSO) cell culture grade (AppliChem) at 10 mg/mL. The stocks of potassium clavulanate (CLA) (Sigma-Aldrich) were prepared in phosphate buffer at 0.1 M, pH 6.0.

**Table 1:** Bacterial strains, plasmids, and cell lines used in this study.

| Bacterial strain, plasmid or cell line | Description | Reference or source |
| --- | --- | --- |
| <u>Bacteria</u> |  |  |
| *Escherichia coli* |  |  |
| JM109 | *recA1 endA1 gyr96 thi hsdR17 supE44 relA1 Δ(lac-proAB) [F' traD36 proAB lac<sup>q</sup>ZΔM15]* | Stratagene |
| BL21(DE3) | *fhuA2 [lon] ompT gal (λ DE3) [dcm] ΔhsdS λ DE3 = λ sBamHlo ΔEcoRI-B int::(lacI::PlacUV5::T7 gene1) i21 Δnin5* | New England Biolabs |
| *Mycobacterium smegmatis* |  |  |
| mc<sup>2</sup> 155 (WT) | High-transformation-efficiency mutant of *M. smegmatis* ATCC 607 | 35 |
| <u>Plasmids</u> |  |  |
| PLJR962 (8881 bp) | Integrative plasmid Sth1 *dcas9* Tet<sup>R</sup> and Kan<sup>R</sup> L5 Int *attP* for *M. smegmatis* | 34 |
| <u>Cell lines</u> |  |  |
| THP-1 | Human-derived monocyte cell line | ATCC, TIB-202 |

### Cloning of the CRISPRi plasmid and preparation of the *cwlM* knockdown mutant

The cloning techniques were carried out as formerly described [6, 34, 36]. The non-template strand of the gene of interest (GOI), *cwlM* (*MSMEG_6935*) was targeted, to produce efficient silencing. Since *cwlM* is putatively essential for *Mtb* viability [30], the template strand of *cwlM* was searched for a PAM associated with medium-to-low repression and accordingly guided sgRNA design (**Table 2**). The CRISPRi backbone PLJR962 was amplified in *E. coli* JM109 and digested as previously described [6]. For sgRNA cloning, two complementary primers were designed (PFwd_sgRNA_cwlM: 5’ **-** GGGAACCCGCCGGTGACGCGTGAGC - 3’; PRv_sgRNA_cwlM: 5’ - AAACGCTCACGCGTCACCGGCGGGT - 3’), synthesized (Eurofins Genomics), annealed, and ligated to the backbone vector with T4 DNA ligase (400 U/µL; NEB). To obtain the interference plasmid, *E. coli* competent cells were transformed with the ligation mixture by double heat-shock, and the resulting recombinant DNA was purified with the NZYMiniprep kit (NZYTech). *M. smegmatis* competent cells were electroporated with 300 ng of recombinant DNA in the presence of 10% glycerol to generate the *cwlM* knockdown mutant (KDM) [6, 37].

**Table 2:**
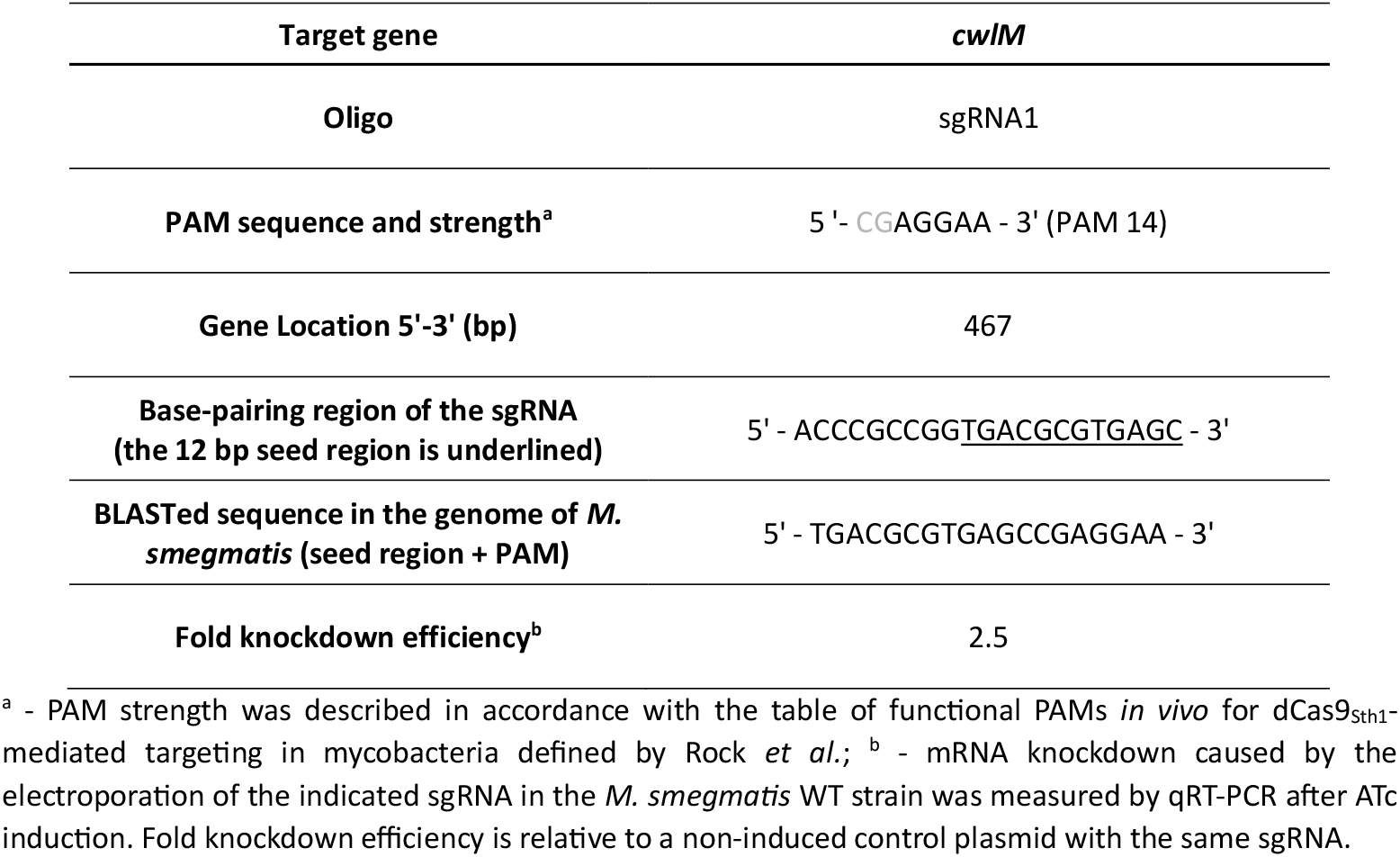
sgRNAs used to target *cwlM* in *M. smegmatis*.

### RNA extraction and quantitative reverse-transcription PCR (qRT-PCR)

The collection of cultures for total RNA extraction was performed as previously described [6]. Briefly, induction with 100 ng/mL of ATc occurred at an optical density at 600 nm (OD_600_) of about 0.1 and the cultures were then allowed to grow for 6 h at 37°C. The cells were harvested, resuspended in 500 µL of RNAprotect Bacteria Reagent (Qiagen) and stored at -80°C. Lysis was carried out with 350 µL of buffer NR (NZYTech) and 3.5 µL of beta-mercaptoethanol (Merck), followed by mechanical disruption in a BeadBug™ (Benchmark Scientific). Bead-beating occurred for 4 cycles (each with 1 min) at maximum speed, with 1 min incubations on ice between each cycle. After, the lysate was transferred into the NZYSpin Homogenization column (NZYTech), and the following steps were performed according to the instructions provided by the NZYTotal RNA Isolation Kit. RNA was eluted with warm DEPC water (Invitrogen) and rigorously treated with TurboDNase (Invitrogen) to prevent genomic DNA contamination. The obtained RNA was quantified using Nanodrop™ and the lack of DNA contamination was corroborated by PCR on the housekeeping gene *sigA*, followed by gel electrophoresis analysis [6].

Reverse transcription reactions were performed using 100 ng of purified RNA as template, following the instructions of the NZY First-Strand cDNA Synthesis kit (NZYTech). Afterwards, cDNA was quantified using the NZYSupreme qPCR Green Master Mix (2x), ROX (NZYTech) on a QuantStudio™ 7 Flex Real-Time PCR System (Applied Biosystems). The 40-cycle qPCR reaction ensued in this way: 95°C for 2 min, then 95°C for 5 s and 60°C for 30 s. The employed primers (Eurofins Genomics; **Table 3**) were designed to amplify a specific product between 100-200 bps, and their amplification efficiency was assessed by calibration curves and proven to be 80-110%. For every run, a minimum of two technical replicates and one negative control were utilized, and the mean C_T_ was calculated. Data were analyzed using the ΔΔC_T_ method with *sigA* (*MSMEG_2758*) as a reference gene and *M. smegmatis* WT as the calibrator sample [34, 36, 38]. Two independent measurements were used to quantify the target mRNA sequences [6, 39].

**Table 3:**
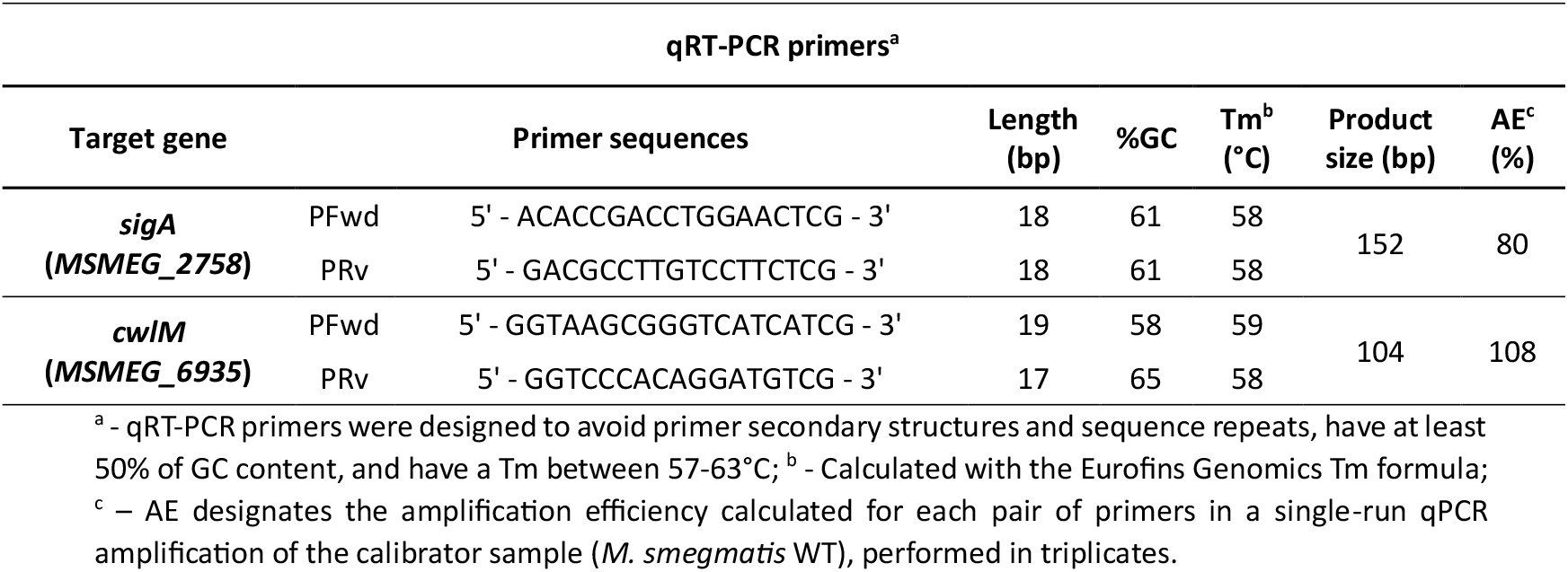
Primers used to quantify the mRNA expression of target genes by qRT-PCR in *M. smegmatis*.

### Phenotyping

To evaluate cell viability differences following CRISPRi induction, spotting assays were performed as previously described [6]. Briefly, cultures of *M. smegmatis* were allowed to reach the log phase of growth, normalized to a theoretical OD_600_ of 0.001, and serially diluted. A 5 µL aliquot from each dilution was plated on agar media, which could be devoid of antibiotics or contain meropenem (MEM; 0.25, 0.5, and 1 µg/mL) or cefotaxime (CTX; 8, 16, and 32 µg/mL) at different concentrations, as indicated in brackets. The plating of each bacterial suspension occurred in both induced and non-induced conditions, following incubation at 37°C for 2 days. At least three independent experiments were performed.

### Minimum inhibitory concentration (MIC) determination

To uncover differences in antibiotic susceptibility, the MICs of three beta-lactams (AMX, CTX and MEM), with and without the beta-lactamase inhibitor clavulanate, as well as one glycopeptide (vancomycin) and two first-line antimycobacterial agents (INH and EMB), were determined as formerly described [6]. The MIC was annotated as the lowest concentration of antibiotic capable of preventing visible bacterial growth. The median MIC values were calculated from three independent experiments.

### Disk diffusion assays

To determine susceptibility differences on solid media following CRISPRi induction, disk diffusion assays were performed with MEM or AMX+CLA, as described previously [21, 40]. Briefly, the cultures of *M. smegmatis* were grown to log phase, normalized to a theoretical OD_600_ of 0.05, and uniformly distributed over the 7H10 (+ ATc) plates using a pour-plate technique. Subsequently, antibiotics were prepared at the indicated test concentrations (MEM: 256 and 512 µg/mL; AMX: 512 and 1024 µg/mL; CLA: 2.5 µg/mL). Then, 20 μL of each antibiotic dilution was added to 9 mm filter paper disks (Filterlab), which were allowed to dry and placed onto the mycobacterial lawn using a sterile needle. The plates were then incubated at 37°C for 2 days, after which photographs were captured on an iBright FL1500 Imaging System (Invitrogen) and the diameter of the zone of inhibition (halo) was measured (in mm) using the ImageJ software. Three independent experiments were performed.

### THP-1 infection assays and intracellular survival evaluation

To explore the role of CwlM in intracellular survival, THP-1-derived macrophages were infected with control and mutant bacteria. THP-1 monocytes were passaged every 3-4 days, being kept at a density above 2 × 10^5^ cells/mL. When necessary, the cells were seeded in 48-well flat-bottom tissue culture plates at a density of 1.5 × 10^5^ cells/well and differentiated into macrophages with 20 nM of phorbol 12-myristate 13-acetate (PMA; Sigma-Aldrich) for 24 h [41]. On the day of THP-1 seeding, the bacterial cultures were diluted to an OD_600_ of about 0.007 and incubated at 37°C with shaking till an OD_600_ of approximately 0.1 was reached. At that point (∼24 h before infection), the cultures were incubated with or without 100 ng/mL of ATc.

Approximately 24 h before infection, the cell media containing PMA was removed and replenished with prewarmed supplemented RPMI media. On the day of infection, bacterial cultures were centrifuged at 3,000 x g for 5 min and washed with 5 mL of PBS 1 X (Gibco) [42]. The resultant pellets were then resuspended in supplemented RPMI media, subjected to an ultrasonic bath for 5 min, and centrifuged at 500 x g for 1 min to obtain a single-cell suspension [43]. After discarding the cell media from the seeding plates, THP-1-derived macrophages were infected with mycobacteria at a multiplicity of infection (MOI) of 5 and, incubated at 37°C with 5% CO_2_ for several timepoints (1 h; 4 h; 24 h) [6,42]. After one hour of internalization, the infection media was discarded, macrophages were washed thrice with prewarmed PBS 1 X, and resuspended in supplemented RPMI media containing 50 µg/mL gentamicin to kill extracellular bacteria and 100 ng/mL of ATc, when suitable. Following distinct pre-determined incubation times (T_1_, T_4_, T_24_), the media was discarded, the cells were washed thrice with PBS 1 X and, incubated at 37°C with a 0.05% IGEPAL solution for 15 min to induce lysis [43]. After, the lysates were serially diluted in sterile water and plated onto 7H10 plates, which were incubated for 2 days at 37°C with 5% CO_2_ [42]. Using an inverted microscope (Nikon TMS Inverted Phase Contrast Microscope), the colony-forming units (CFUs) of three independent biological replicates, each counting three technical replicates, were counted.

### Cloning of recombinant CwlM_TB_

To overexpress and characterize the CwlM_TB_ protein, the *cwlM* gene (*Rv3915*) was cloned in frame with a C-terminal polyhistidine (His6) tag into the pET29b backbone, as previously described [40, 44]. The oligonucleotides PFwd_*cwlM* (5’ – GACCATATGCCGAGTCCGCGCCGCG – 3’) and PRv_*cwlM* (5’ – GCCCTCGAGAGAACCGCCGAGTCTACC – 3’) were used to amplify the *cwlM* gene from 30 ng of *Mtb* H37Rv genomic DNA using 0.025 U/µL of Supreme NZYProof DNAPolymerase (NZYTech). The PCR was conducted on a thermocycler (Life Technology - SimpliAmp), with the following PCR program: 96°C for 4 min, 1 cycle; 96°C for 30 s, 63°C for 30 s and 72°C for 40 s, 30 cycles; 72°C for 10 min, 1 cycle. The 1234 bp PCR product was confirmed by 0.8% agarose gel electrophoresis and visualized using the iBright™ FL1500. Afterwards, the PCR reaction was scaled up, the PCR product was purified using the NZYGelpure kit (NZYTech) and, 800 ng of purified PCR product and 600 ng of the pET29b plasmid were then digested at 37°C for 3 h with 2.5 µL of FastDigest Xho*I* and Nde*I* (ThermoScientific) in a final volume of 80 µL. The reaction products were purified using the NZYGelpure kit (NZYTech) and separated by a 0.8% agarose gel electrophoresis, followed by visualization on iBright™ FL1500. The digested fragment was then ligated to 50 ng of digested pET29b with 80 U of T4 DNA ligase (400 U/µL; NEB) using a vector-to-insert ratio of 1:5, at 22°C for 1 h in a ThermoMixer C (Eppendorf). Afterwards, *E. coli* JM109 chemically competent cells were transformed with 5 μL of ligation mixture using a double heat-shock approach, and the recombinant DNA was purified according to the instructions of the NZYMiniprep kit (NZYTech). The obtained recombinant plasmid was then quantified, evaluated for purity, and sequenced using primers that specifically amplify the T7 RNA polymerase promoter PFwd_T7 (5’ – TAATACGACTCACTATAGGG – 3’) and terminator PRv_T7 (5’ – CTAGTTATTGCTCAGCGG – 3’).

### Expression of recombinant CwlM_TB_ in *E. coli* BL21(DE3) and SDS-PAGE analysis

*E. coli* BL21 (DE3) chemically competent cells were transformed with 300 ng of the pET29b:*cwlM* construct using a double-heat shock procedure [40]. Inoculums from *E. coli* BL21(DE3):pET29b:*cwlM* transformants were prepared and incubated at 37°C with shaking until the OD_600_ was about 0.5. Then, cultures were induced with 1 μM of IPTG (NZYTech), incubated in two distinct conditions, at 37°C for 3 h and at 16°C overnight, collected by centrifugation, and frozen at -20°C [40]. The next day, pellets were resuspended in a lysis buffer solution [(50 mM Tris, 300 mM NaCl), supplemented with 1% of a cocktail of protease inhibitors (Calbiochem), 10 μg/mL DNAse, 10% glycerol and 0.1% Triton X-100], sonicated twice for 60 seconds (0.5 cycles, 100% amplitude, pulse on, on ice), centrifuged at 8,000 x g, 4°C for 20 min, separated from the supernatant, and stored at -20°C. SDS-PAGE analysis of the obtained pellets and supernatants was performed in a Mini-PROTEAN® Tetra Cell to verify the presence of the protein of interest, according to the standard protocol Hand Casting Polyacrylamide Gels by Bio-Rad [40]. To prepare the ready-to-load samples, 6 μL of loading dye (5 X SDS-PAGE Sample Loading Buffer; NZYTech) were combined with 24 μL of the supernatants. Pellets were resuspended in 24 μL of running buffer 1 X (50 mM Tris, 500 mM glycine (pH 8.3), 0.1% w/v SDS), and then mixed with 6 μL of loading dye (5 X SDS-PAGE Sample Loading Buffer). Afterwards, the samples were heated at 100°C for 5 min and iced for 1 min. An amount of 20 μL of each sample was applied to the gel, the stacking gel ran at 80 V and the resolving gel ran at 120 V. After the run, the gel was stained with BlueSafe (NZYTech) for approximately 30 min. Based on the comparison of soluble protein yields under both induction conditions, we chose to induce 2 L of culture at 16°C overnight [40].

### Purification of CwlM_TB_

After induction, the scaled-up 2 L culture was collected by centrifugation, and the resulting pellet of *E. coli* BL21(DE3) cells expressing CwlM_TB_ was thawed on ice and resuspended in 40 mL of lysis buffer [40]. Lysis was achieved by sonication in 10 cycles of 30 s, 50% amplitude, interpolated by 40 s rest periods. Afterwards, the lysate was centrifuged (Centrifuge 5810 - Eppendorf) at 11,200 x g, 4°C, for 45 min and the supernatant was filtered using 0.45 μM and 0.2 μM filters (GE Healthcare). The soluble extract was then loaded in ÄKTAprime plus and purification of the His_6_-tagged CwlM_TB_ protein was carried out with a HisTrap FF Crude 1 mL Column (Cytiva), according to the instructions provided by the manufacturer. Washing buffer (50 mM Tris, 300 mM NaCl, 40 mM Imidazole, 1 mM DTT, pH 8.0), elution buffer (50 mM Tris, 300 mM NaCl, 300 mM Imidazole, 1 mM DTT, pH 8.0), MilliQ water, and 20% ethanol were also loaded in the system when appropriate. The bound CwlM_TB_ protein was eluted in the presence of imidazole, a competitive agent of His_6_-tagged proteins [45]. After elution, all fractions were collected, pooled, and the buffer was exchanged by overnight dialysis at 4°C with magnetic stirring, using dialysis buffer (20 mM Tris, 150 mM NaCl, 25% glycerol, 1 mM DTT) [40]. Next, the protein sample was centrifuged at 10,000 x g, 4°C, for 10 min and filtered using 0.2 μM filters. The purified CwlM_TB_ protein sample was subsequently quantified by the Bradford method, using bovine serum albumin (BSA, 10 μg/mL) (Sigma-Aldrich) as a standard [40].

### Zymography study to characterize the enzymatic activity of the CwlM_TB_ protein

To uncover whether the CwlM_TB_ protein has PG-hydrolytic activity, a zymography assay was carried out against PG obtained from *M. luteus*. First, the bacteria were grown at 37°C for two days until exhaustion, centrifuged at 4,000 x g, 4°C, for 10 min, and the pellet was washed once with MilliQ water. The pellet was then resuspended in MilliQ water, autoclaved, and harvested by centrifugation at 12,000 x g, 4°C for 20 min. After discarding the supernatant, the pellet was dried at 37°C for 24 h, weighed, and a 2% solution of *M. luteus* PG in MilliQ water was prepared and stored at -20°C. The zymogram gel was prepared according to Bio-Rad instructions (12% running gel and 5% stacking gel) and the 2% solution of *M. luteus* PG was added to the appropriate volume of water to achieve a final concentration of 0.2% [40]. An equivalent SDS-PAGE gel without *M. luteus* PG was also prepared to track the run. Loaded samples comprised 5 μg of BSA (negative control, NZYTech), 5-10 μg of CwlM_TB_ and 1-2 μg of Lysozyme (Lys, positive control, Sigma-Aldrich) in sample buffer (62.5 mM Tris-HCl, pH 6.8, 2% SDS, 5% β-mercaptoethanol, 20% glycerol, 0.01% bromophenol blue). Samples were heated at 100°C for 5 min and iced for 1 min. A small volume (4 μL) of the NZYColour Protein Marker II (NZYTech) and 20 μL of each sample were loaded onto the gels, which were subsequently run and stained with BlueSafe. The zymogram gel was washed in distilled water for 30 min at RT, and then incubated in renaturation buffer (5% Triton X-100, 25 mM Tris-HCl, pH 7.5) at 37°C, for 16 h, with shaking at 120 rpm. After renaturation, the gel was washed twice for 15 min, tinted with a staining solution (0.2% methylene blue, 0.01% KOH) for 30 min, and repeatedly de-stained with distilled water until the protein ladder was visible 40]. Gel visualization was performed using the iBright™ FL1500. The zymography assay was only performed once.

### Statistical analysis

GraphPad (version 8.4.3.686) was used for statistical analysis and data are here presented as mean ± SEM, unless otherwise specified. The qRT-PCR fold expression values were tested for normality using Q-Q plots, whereas the CFUs counts were analyzed through the Shapiro-Wilk Test. Multiple group comparisons were accomplished using ordinary one-way ANOVA and pairwise comparisons of selected groups were evaluated via the Holm-Sidak test. All the prerequisites of the tests were verified.

## Results

### Construction and characterization of the *cwlM* knockdown mutant

The dCas9_Sth1_-based CRISPRi system [34] was implemented to construct a *cwlM* KDM in *M. smegmatis*. To produce an efficient repression of *cwlM* (*MSMEG_6935*), the non-template strand was targeted. The efficiency of CRISPRi-mediated knockdown is influenced by several factors, including PAM strength, sgRNA-DNA_target_ complementarity, target site, and GC content [6, 34]. The strength of the employed PAM is often directly associated with the produced knockdown level and the resultant growth inhibition phenotypes [6]. Therefore, the repression of essential genes mediated by strong PAMs might cause cell death. Since *cwlM* is allegedly essential to the viability of *Mtb* [30], the first half of the template strand of the gene was searched for a PAM associated with medium-to-low repression (**Figure 1a**). Choosing a target site in the first half of the GOI is important to ensure that the produced mRNAs do not originate truncated proteins retaining some degree of activity. Although polar effects are common when CRISPRi is employed [34, 38-39], *cwlM* is a non-operonic gene and its repression is not likely to affect the transcription of neighbouring genes. Before mycobacterial transformation, the *cwlM*-targeting CRISPRi plasmid was verified by Sanger Sequencing (**Additional file 1: Figure S1**).

**Figure 1.**
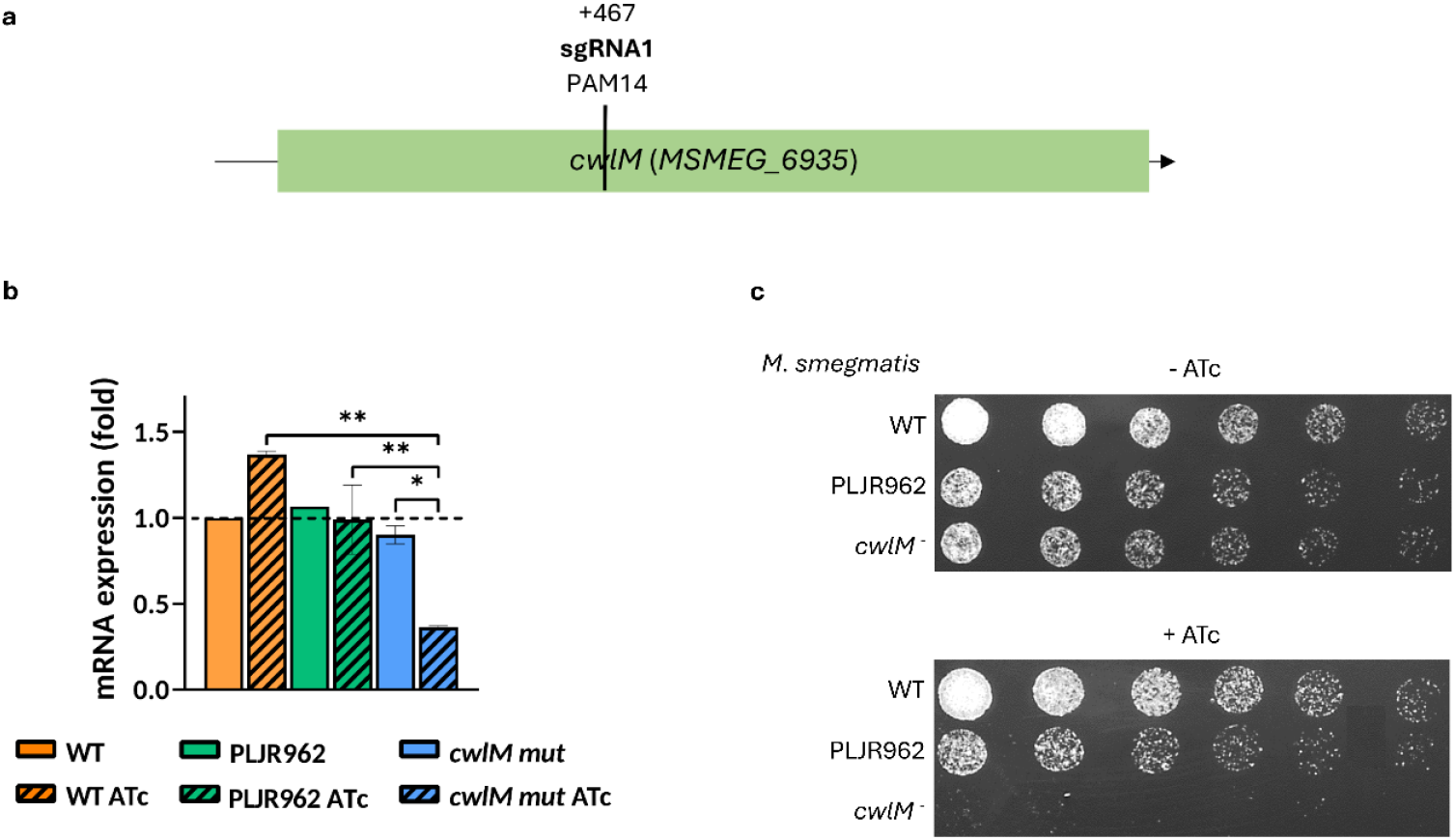
CRISPRi-mediated *cwlM* silencing in *M. smegmatis* and resulting phenotypical and mRNA expression differences. **(a)** sgRNA-mediated repression of the *cwlM* gene in *M. smegmatis*, detailing target site (+467 bps) and the employed PAM strength (PAM14). **(b)** Relative gene expression of *cwlM* normalized to *sigA*, with (stripped bars) and without (smooth bars) 100 ng/mL of ATc treatment for 6 h, evaluated through qRT-PCR assays (n=2). PLJR962 is a non-targeting empty vector control. The mRNA expression levels of the calibrator sample (*M. smegmatis* WT) are shown as dashed lines. The error bars represent the standard error of the mean (SEM). Comparisons between multiple groups were made using one-way ANOVA, with significance levels: * *P* < 0.05; ** *P* < 0.01. **(c)** Spotting dilution assays of the negative controls (*M. smegmatis* WT, PLJR962) and of the *cwlM* knockdown mutant (n=3).

Following this, the normalized relative gene expression of *cwlM* was determined for the controls *M. smegmatis* WT and PLJR962 and the *cwlM* KDM, with and without inducer, at 6 h post-induction. The results show that the *cwlM* gene was successfully silenced (**Figure 1b**). The sgRNA1-mediated knockdown of *cwlM* produced a very significant decrease in the target mRNA expression levels compared to WT ATc (3.8-fold; *P* = 0.0015) and to PLJR962 ATc (2.5-fold; *P* = 0.0029). The repression of *cwlM* was also determined as significant relative to the respective non-induced control (2.7-fold; *P* = 0.02) (**Figure 1b**).

Afterwards, spotting dilution assays were performed to facilitate the phenotypical characterization of the *cwlM* KDM. The growth of negative controls *M. smegmatis* WT and PLJR962 was not affected by ATc treatment (**Figure 1c**). Conversely, the *cwlM* KDM suffered a severe reduction in cell viability in the presence of inducer (**Figure 1c**).

### Influence of CwlM depletion on antibiotic susceptibility

To uncover whether the depletion of CwlM provokes any considerable changes in antibiotic susceptibility, the MICs of EMB, INH, VAN and of three beta-lactams (AMX, CTX and MEM) in the presence or absence of clavulanate, were determined against the control strains and the *cwlM* KDM (**Figure 2**). As expected, *M. smegmatis* WT and PLJR962 did not suffer extensive changes in susceptibility to the tested antibiotics, even with inducer [40]. Notably, the knockdown of *cwlM* promoted a subtle increase in susceptibility to all tested antibiotics. Consistent with commonly accepted thresholds for microdilution susceptibility studies, 2-fold MIC changes were attributed to technical variability; therefore, only MIC differences of ≥ 4-fold were considered biologically meaningful. The depletion of CwlM did not seem to substantially affect the susceptibility of mycobacteria to first-line agents EMB and INH, AMX (+CLA), MEM (+CLA) and to the PG cross-linking inhibitor VAN [40]. Conversely, the repression of *cwlM* promoted a noteworthy increase in susceptibility (≥ 4-fold) to CTX (+CLA) [40]. The addition of CLA to AMX and CTX produced 16-fold and 2-fold MIC reductions, respectively. On the other hand, the addition of CLA to MEM did not provoke any MIC changes.

**Figure 2.**
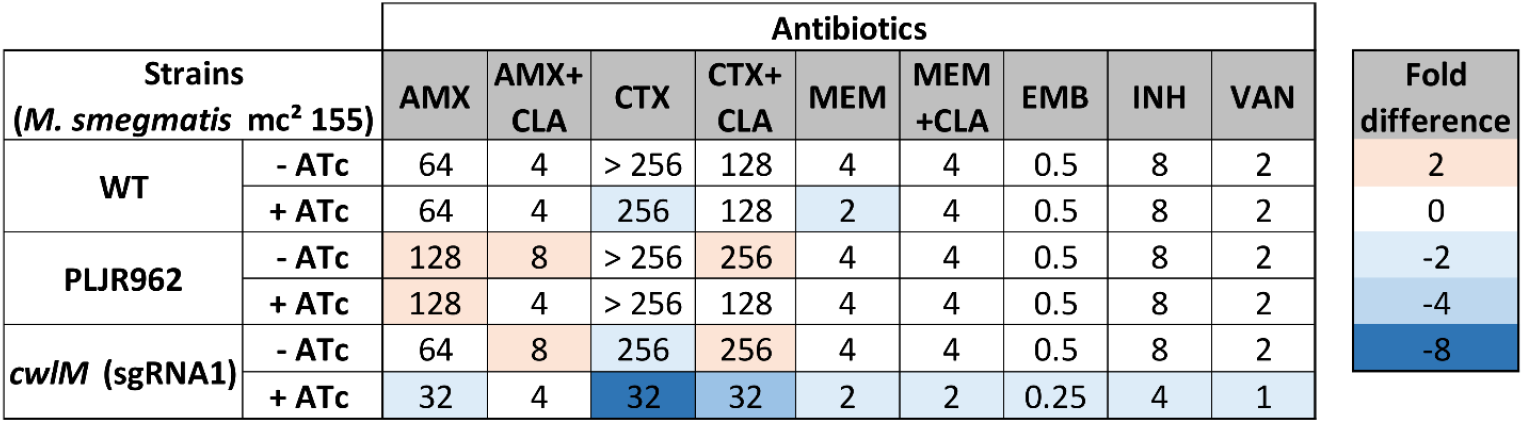
Antibiotic susceptibility assays of the negative controls and the *cwlM* knockdown mutant. Heatmap of the fold differences in the minimum inhibitory concentrations (MICs; in μg/mL) between the *cwlM* knockdown mutant, the non-targeting control PLJR962 and *M. smegmatis* WT, with and without 100 ng/mL of ATc. The red shades depict an increase in the MIC while the blue shades depict a decrease in the MIC, when compared to the WT strain. AMX, amoxicillin; CLA, clavulanate; CTX, cefotaxime; EMB, ethambutol; INH, isoniazid; MEM, meropenem; VAN, vancomycin.

### CwlM knockdown promotes increased susceptibility to beta-lactams on solid media

To validate these observations, spotting dilution assays of the control *M. smegmatis* WT and the *cwlM* KDM were performed in the presence of varying concentrations of CTX and MEM, with and without inducer (**Figure 3a**). As formerly demonstrated, the *cwlM* KDM suffered a severe growth defect in the presence of 100 ng/mL of inducer. The viability of *M. smegmatis* WT remained mostly unaffected regardless of the concentration of antibiotics tested. Nevertheless, a marked viability defect was observed with 1 μg/mL of MEM [40]. Moreover, the growth profile of the *cwlM* KDM in non-induced conditions was very similar to that of the WT strain, with both antibiotics. On the other hand, the induced *cwlM* KDM presented a slight viability loss with both MEM and CTX when compared to the control strain [40]. In fact, the ability of the induced mutant to form a spot was indirectly proportional to the concentration of antibiotic used.

**Figure 3.**
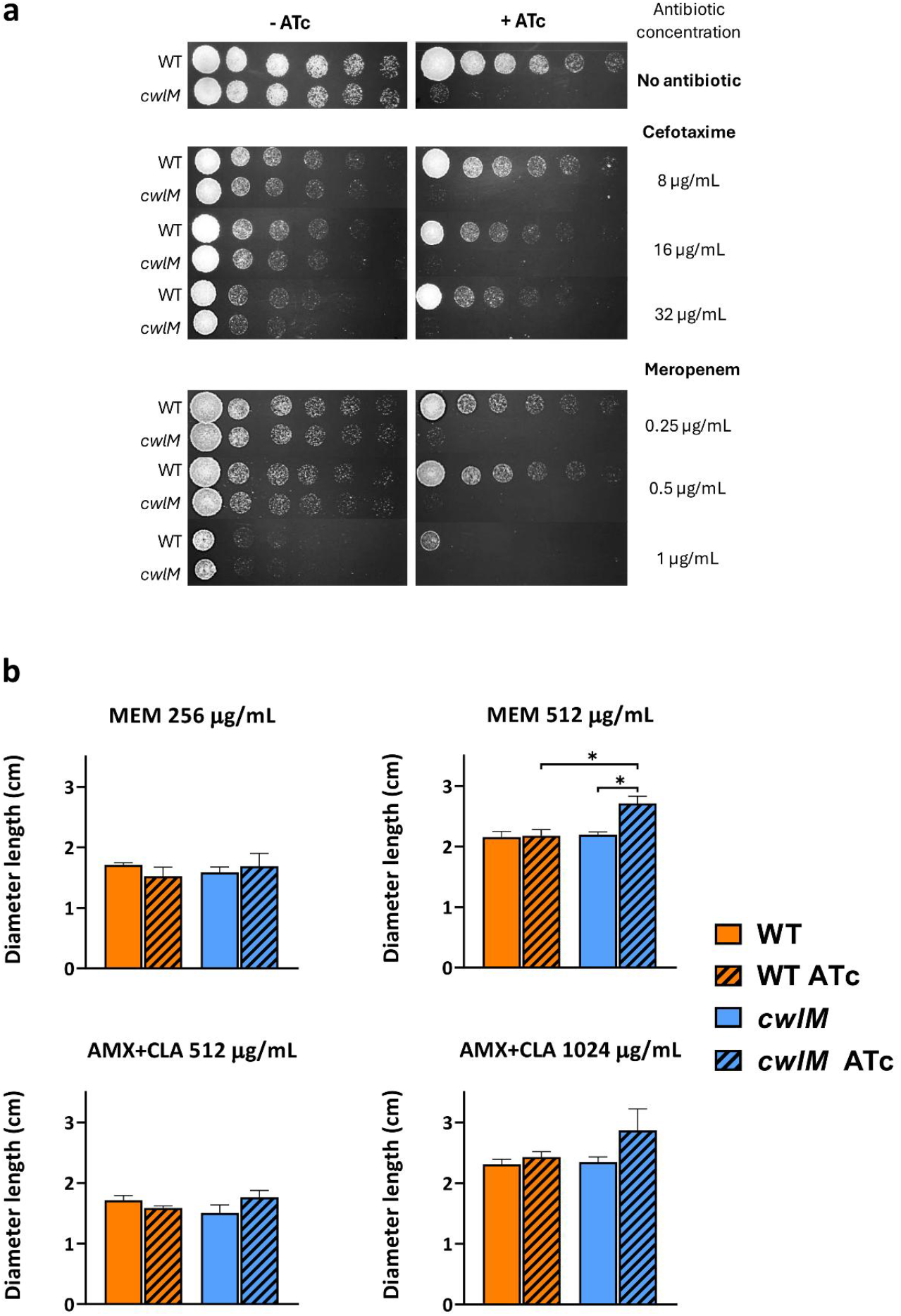
Susceptibility assays of the negative controls and the *cwlM* knockdown mutant on solid media. **(a)** Spotting dilution assays of the negative control strain *M. smegmatis* WT and the *cwlM* knockdown mutant, with and without 100 ng/mL of ATc, without any antibiotics and in presence of varying concentrations of cefotaxime (8, 16, and 32 μg/mL) and of meropenem (0.25, 0.5, and 1 μg/mL). **(b)** Average length of the diameter of the inhibition halo (in cm) obtained for amoxicillin supplemented with clavulanate (AMX+CLA) and meropenem (MEM) antibiotic disk assays performed with the negative control *M. smegmatis* WT and the *cwlM* knockdown mutant, with (striped bars) and without (smooth bars) 100 ng/mL of ATc (n=3). The diameter of the inhibition halo was measured using ImageJ. The error bars correspond to the standard deviation. Multiple group comparisons were calculated using ordinary one-way ANOVA followed by the Holm-Sidak’s test (* *P* < 0.05).

To further consolidate these findings, AMX+CLA and MEM disk diffusion assays were performed for *M. smegmatis* WT and the *cwlM* KDM, with and devoid of inducer (**Figure 3b**). As follows, the concentrations of antibiotics used to produce a visible halo effect had to be optimized. For AMX+CLA, this concentration was 8-fold greater than the MIC determined in liquid media and for MEM, it was 64-fold the MIC. CTX was excluded from this assay due to the necessity of preparing a highly concentrated stock solution defying solubility limitations. With 256 µg/mL of MEM, no significant differences in the halo diameter were observed between the strains, regardless of ATc addition. With 512 µg/mL of MEM, the induced *cwlM* KDM displayed a significant increase in susceptibility to the carbapenem when compared to the induced WT strain (1.25-fold; *P* = 0.012) and to the respective non-induced control (1.24-fold; *P* = 0.015) [40]. With 512 µg/mL of AMX+CLA, no significant differences in the halo diameter were observed between the WT strain and the *cwlM* KDM, independently of ATc addition. With 1024 µg/mL of AMX+CLA, the induced *cwlM* KDM displayed a modest increase in susceptibility to the penicillin, albeit without significance [40].

### Impact of CwlM depletion in the killing of mycobacteria within THP-1-derived macrophages

To uncover whether CwlM contributes to survival within host macrophages, THP-1-derived macrophages were infected with the control strains and the *cwlM* KDM at an MOI of 5. The intracellular survival (in CFUs/mL) was then assessed following cell lysis at 1 h, 4 h and 24 h post-infection. No considerable differences in *in vitro* growth kinetics were observed among the study strains prior to infection; OD_600_ values differed by less than 0.23 across all strains (**Additional file 1: Table S1**).

As expected, the viability of mycobacteria reduced over time [6]. Moreover, the mean intracellular survival of the negative controls *M. smegmatis* WT and PLJR962 was not significantly affected by the presence of inducer. The knockdown of *cwlM* induced a very significant reduction in intracellular survival at all time points (*P* < 0.001, when compared to *M. smegmatis* WT ATc, PLJR962 ATc and the corresponding non-induced control) (**Figure 4**). Notably, the survival of the induced *cwlM* knockdown mutant within host macrophages consistently decreased over time, being significantly lower than that of *M. smegmatis* PLJR962 ATc (3.1-fold; *P* = 0.024) and of the respective non-induced control (6.3-fold; *P* < 0.001) at T_24_ (**Additional file 1: Figure S2**). These results indicate that the depletion of CwlM might facilitate the macrophage-mediated immune recognition of mycobacteria, therefore promoting a faster clearance of intracellular mycobacteria.

**Figure 4.**
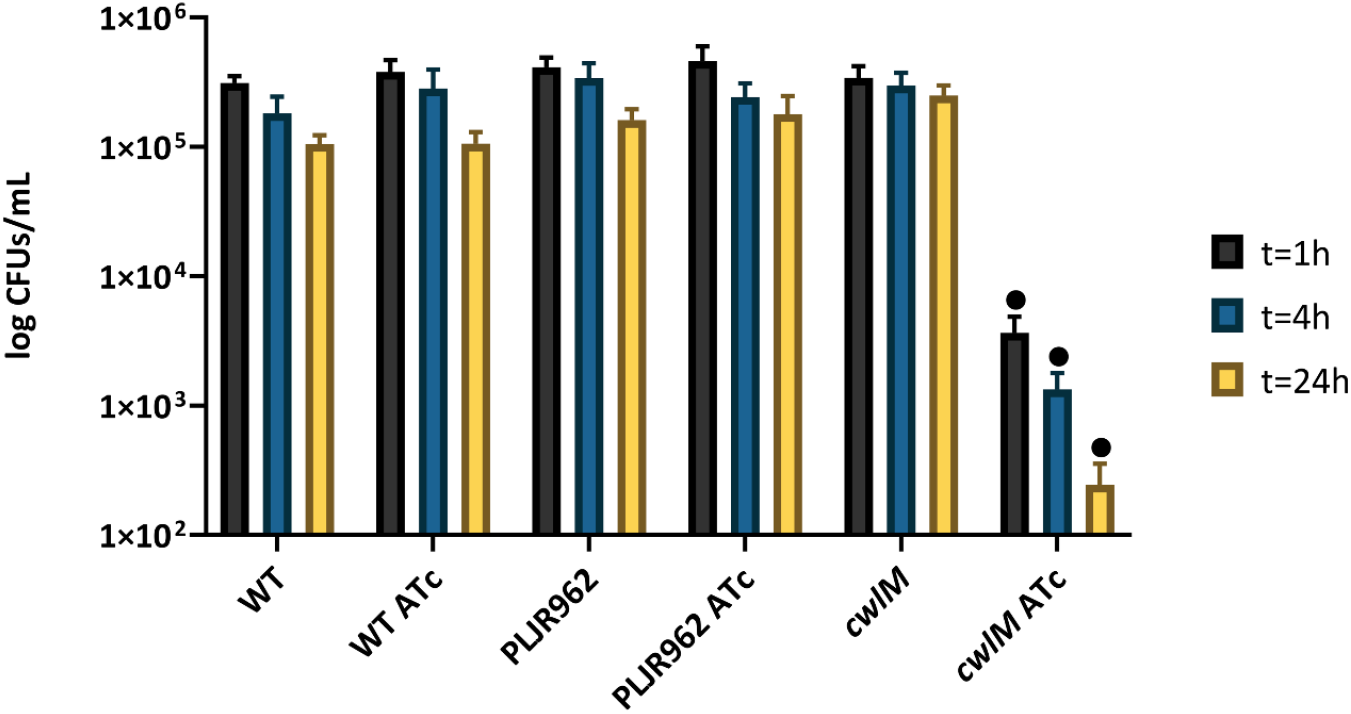
Mean survival, in CFUs/mL, of bacteria recovered from infected and disrupted PMA-differentiated THP-1-derived macrophages. (n=3). The disruption macrophages occurred at 1 h, 4 h, and 24 h post-infection. Comparative effect with and without 100 ng/mL of inducer (ATc). Error bars represent the standard error of the mean (SEM). Multiple comparisons were calculated using one-way ANOVA (• *P* < 0.001 relatively to all pre-selected control strains).

### Cloning, expression and purification of recombinant CwlM_TB_

To produce and characterize the CwlM_TB_ protein, the pET system was employed to clone the *cwlM* gene (*Rv3915*) in frame with a C-terminal polyhistidine (His_6_) tag [40]. Following the verification of construct integrity through Sanger sequencing analysis (**Additional file 1: Figure S3**), recombinant *E. coli* BL21(DE3) cells were used to overexpress *cwlM* [40]. Induction with 1 µM of IPTG was tested under distinct conditions (37°C for 3 h or at 16°C overnight) to optimize overexpression of the CwlM_TB_ protein (43.9 kDa) in pET29b. SDS-PAGE analysis of the resulting fractions showed a discernible band with the expected size of 43.9 kDa in the fourth lane (CwlM at 16°C overnight), compared to the non-induced (N.I) control (**Additional file 1: Figure S4a**) [40]. Overall, these results demonstrate that CwlM_TB_ is only effectively produced when the cells are grown at 16°C overnight. Based on these findings, a scale-up culture of 2 L of *E. coli* BL21(DE3):pET-29b:*cwlM* was prepared, grown and induced under optimal expression conditions. SDS-PAGE analysis of the resulting fractions confirmed the successful production of CwlM_TB_, with a visible band at 43.9 kDa in the third lane, which was absent in the N.I. control (**Additional file 1: Figure S4b**) [40].

Purification of CwlM_TB_ yielded a strong absorbance peak between fractions 7 and 11, with a maximum of about 250 mAμ at 280 nm (**Figure 5**). SDS-PAGE analysis confirmed that fractions 6-9 contained most of the concentrated protein, as these fractions exhibited large bands at ∼43.9 kDa in the gel, matching the expected molecular weight of CwlM_TB_ (**Figure 5**). After buffer exchange and imidazole removal, the final purified CwlM_TB_ protein was obtained at a concentration of about 800 μg/μL [40]. SDS-PAGE analysis following dialysis (**Figure 5**) revealed several bands: while the inferior bands might indicate a trivial amount of degraded protein, the upper band might represent an aggregate of CwlM with a smaller fragment.

**Figure 5.**
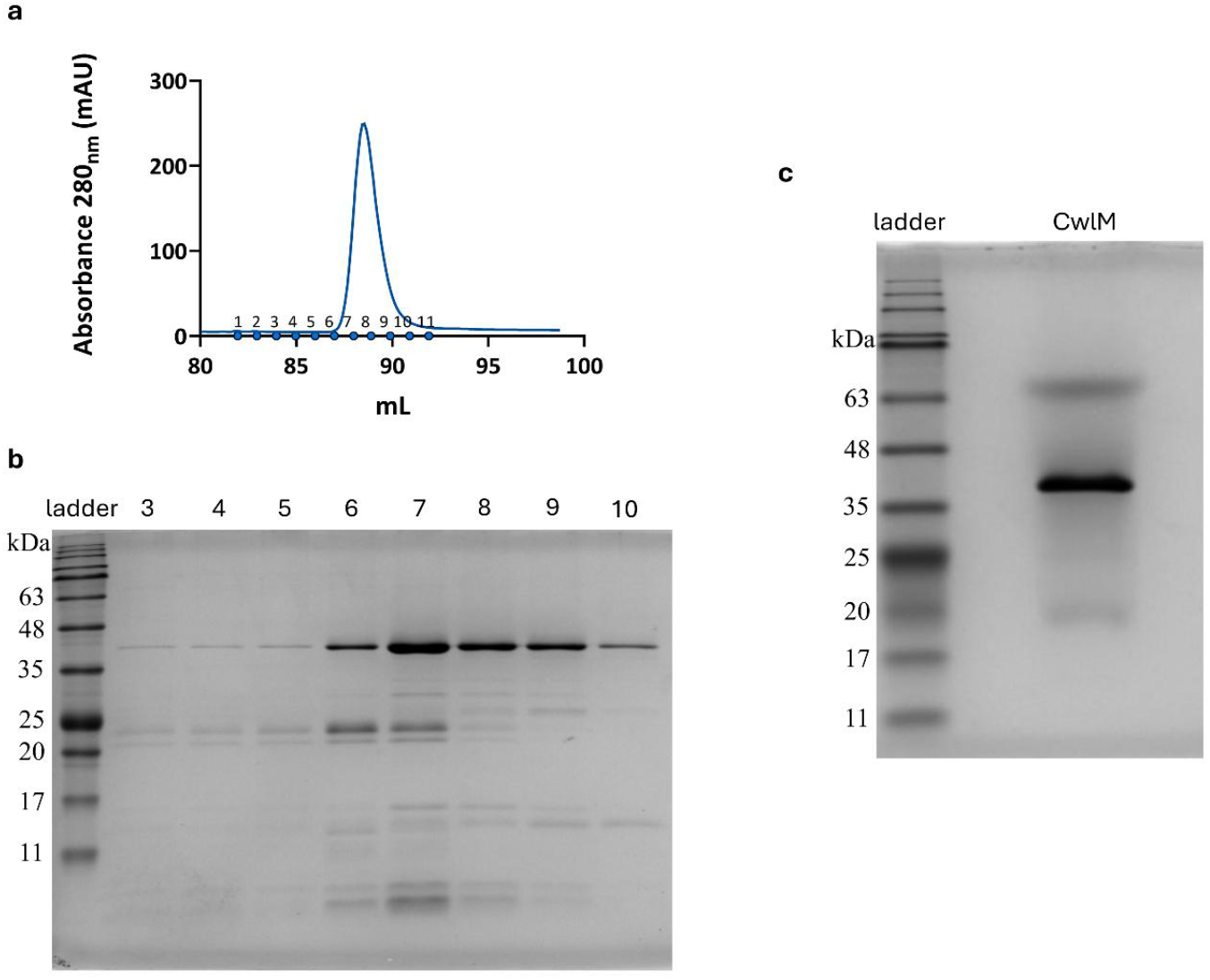
Purification of the CwlM protein. **(a)** Absorbance at 280 nm measured by the ÄKTAprime plus system during the elution step of the affinity purification of CwlM from an induced filtrate of *E. coli* BL21(DE3):pET-29b:*cwlM*. The collected fractions are 1 through 11 and the last dot is waste. **(b)** SDS-PAGE gel of the collected fractions of the CwlM protein after purification in the AKTAprime system. Lane order: NZYColour Protein Marker II ladder; fractions 3, 4, 5, 6, 7, 8, 9 and 10. **(c)** SDS-PAGE of the final purified CwlM protein. Lane order: NZYColour Protein Marker II ladder; CwlM protein (predicted size of 43.9 kDa).

### Characterization of enzymatic activity via zymography and plate assays

To elucidate whether CwlM possesses any PG-hydrolytic activity, a zymogram protocol similar to the one described in [30], was performed using PG from *M. luteus* as substrate (**Figure 6a**,**b**). The zymogram displayed one translucent zone around 14 kDa, consistent with the lytic activity of lysozyme over the PG (positive control -Lys) (**Figure 6b**). Nevertheless, the Lys band displays a molecular weight between 17-25 kDa, having shifted from its predicted size. This could have occurred because of the low concentration of buffers in the zymogram gel, which may impact protein migration, or due to partial renaturation of the Lys protein resulting from prolonged heat exposure during the assay. On the other hand, no translucent zone was observed with the negative control BSA (66 kDa), as it lacks PG-hydrolytic activity. When compared to lysozyme, CwlM_TB_ does not appear to possess any hydrolase activity against *M. luteus* PG (**Figure 6b**) [40]. To further validate these results, a plate assay was performed. While the PG-hydrolytic activity of Lys produced a discernible halo on a *M. luteus* lawn, no halo was observable with CwlM_TB_ (**Figure 6c**) [40].

**Figure 6.**
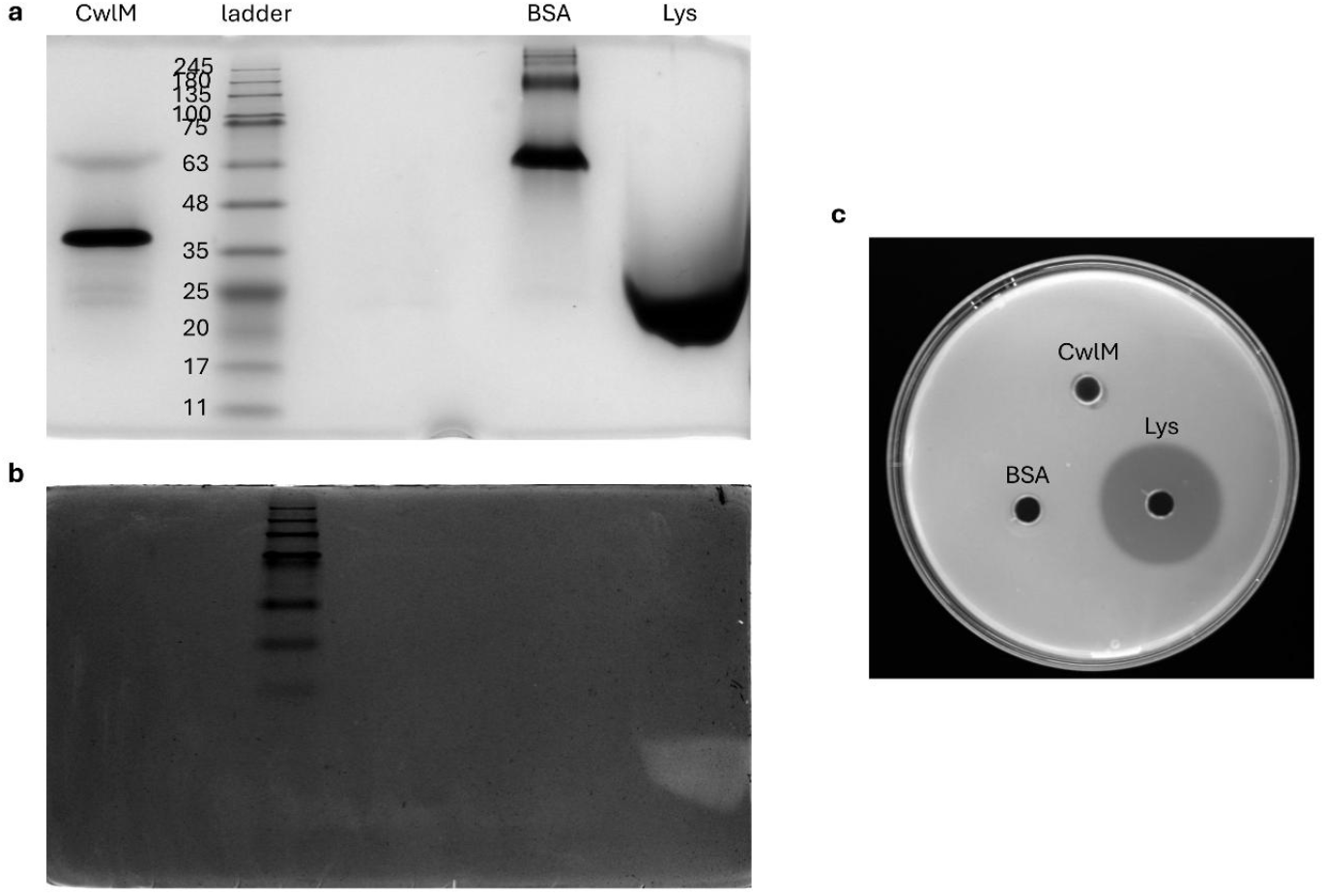
Deciphering the enzymatic activity of CwlM through a zymogram containing 2% of *M. luteus*-derived peptidoglycan. **(a)** SDS-PAGE analysis of CwlM and two controls (BSA and lysozyme). **(b)** The zymogram gel shows that the positive control lysozyme can cleave PG from *M. luteus* while the predicted PG hydrolase CwlM cannot. **(a, b)** Lane order of both gels: CwlM; NZYColour Protein Marker II ladder; BSA (negative control); and Lysozyme (Lys – positive control). **(c)** Plate of *M. luteus* to which the control proteins and CwlM were added. This assay demonstrates the PG-hydrolytic activity of the positive control lysozyme while reiterating the lack of PG-hydrolytic activity of CwlM.

## Discussion

The development of novel antitubercular agents is essential to gain control of DR-TB [2]. The discovery of novel therapeutic targets can be propelled by research into mycobacterial PG biosynthesis enzymes and inhibitors. Although traditionally excluded from DR-TB therapeutic programs, beta-lactams have resurged as promising additions to pipeline drugs, especially when paired with beta-lactamase inhibitors. The WHO now includes two carbapenems in Group C of the medicines recommended for longer MDR-TB therapeutic programs [46]. Recently, we identified several genomic drivers underlying differential beta-lactam susceptibility phenotypes in *Mtb* [29, 47]. Among these, the 4403900 snp A>G in *cwlM* was often associated with increased susceptibility to amoxicillin [29]. Therefore, we sought to investigate the role of CwlM in beta-lactam susceptibility and in intracellular survival using *M. smegmatis* as a model. Due to previous conflicting information, we also purified recombinant CwlM_TB_ to characterize its function. Our findings, discussed below, suggest that CwlM_SMEG_ contributes to beta-lactam resistance and intracellular survival, while CwlM_TB_ lacks PG-hydrolytic activity and might act as a PG biosynthesis regulator instead [31].

First, the qRT-PCR results confirmed the successful CRISPRi-mediated knockdown of *cwlM*. There was barely any evidence of leaky expression phenomena except for a slight reduction in the mRNA levels of *cwlM* observed with the non-induced *cwlM* control. Notably, the observed levels of repression are mild compared to the ∼38.5-fold repression previously reported for target genes in *M. smegmatis* using the same PAM [34]. However, these repression levels were attained with twice the induction period here implemented. Globally, the obtained results demonstrate the efficiency of the dCas9_Sth1_-based CRISPRi system leading to effective target mRNA inhibition and, consequently, affecting the resultant phenotypes.

Consistent with previous reports, the spotting dilution assays revealed a severe growth impairment upon *cwlM* knockdown, further supporting its reported essentiality for mycobacterial survival [30, 48-50]. Notably, even when using a low-strength PAM (PAM14), CRISPRi-mediated silencing of *cwlM* resulted in a near-lethal phenotype, thus underscoring the high vulnerability of *cwlM*, a main criterion for target-based drug discovery [51]. The CwlM protein comprises two PG-binding domains and one MurNAc-LAA domain, with predicted N-acetylmuramoyl-L-alanine amidase activity. A previous report found that the overexpression of CwlM did not result in lysis, presumably due to the lack of two essential catalytic residues (R199, R270) [31]. Instead, the authors proposed a regulatory function for CwlM, demonstrating that phosphorylated CwlM stimulated the activity of MurA and promoted PG biosynthesis. As follows, the depletion of CwlM may well impair PG biosynthesis [32], severely compromising cell division. Hence, the regulatory role of CwlM might affect several pathways and justify its essentiality to cell division and survival.

The MIC assays demonstrated that silencing *cwlM* provoked a slight increase in susceptibility to the tested antibiotics, including beta-lactams. Likewise, previous reports showed that the repression of *cwlM* in *Mycobacterium abscessus* promoted increased susceptibility to beta-lactams [52, 53]. To cease PG biosynthesis and eradicate mycobacteria, all transpeptidases and even other PG-modifying enzymes ought to be inhibited [23]. In our experiments, no major susceptibility differences were observed with AMX, possibly because this penicillin is only able to target PBPs [53]. Whereas the addition of CLA did not greatly impact the MICs of CTX and MEM, it provoked a remarkable 16-fold reduction in the MIC of AMX. This observation is justified by the higher affinity of BlaS for penicillins, as opposed to cephalosporins and carbapenems [55]. In fact, AMX+CLA has been shown to possess bactericidal activity against *Mtb in vitro* [26, 56] and to provide clinical benefit in DR-TB patients [57]. Nevertheless, the WHO has stated that AMX+CLA should not be used without imipenem–cilastatin or meropenem for MDR/RR-TB treatment [46].

Overall, MEM was very efficient at inhibiting mycobacterial growth, compared to AMX and CTX. Although MEM and CTX inhibit both PBPs and Ldts [58], only MEM can do so efficiently and simultaneously escape the hydrolyzing activity of beta-lactamases [59]. Since the activity of MEM is sufficient to inhibit all transpeptidases and, consequently, halt PG biosynthesis, the depletion of CwlM did not greatly affect the MIC of MEM. These observations support the introduction of MEM-containing therapeutic regimens for MDR-TB. Considerable susceptibility differences were only observed with CTX (+CLA), thus suggesting that the activity of CTX is facilitated by the depletion of CwlM, which is expected to cause abnormal cell elongation owing to uncoordinated synthesis of CW layers [31]. We hypothesize that some PG biosynthesis processes may remain active when *cwlM* is repressed and that CTX-mediated inhibition of PG cross-linking is considerably facilitated due to low PG integrity and further accentuates PG polymerization defects.

Similarly to beta-lactams, the glycopeptide VAN can inhibit transpeptidation reactions by irreversibly binding the D-Ala-D-Ala dipeptide [60]. We thus expected the induced *cwlM* KDM to display increased susceptibility to VAN; but observed no such effect. It could be that the D-Ala-D-Ala moiety is protected by the lipid-rich CW of mycobacteria. Still, VAN might be able to inhibit mycobacterial PG biosynthesis when combined with first-line agents EMB and INH [40, 60].

The spotting dilution assays in the presence of varying concentrations of MEM and CTX showed that the depletion of CwlM provokes subtle increases in susceptibility to both beta-lactams, with greater differences observed with CTX, consistent with the MIC assays. In contrast, MEM affected the viability of both *M*. *smegmatis* WT and the induced *cwlM* KDM. The disk diffusion results suggest that a reduction in CwlM protein levels may slightly increase susceptibility to MEM and AMX+CLA, although the obtained fold changes were not biologically relevant, and no statistical significance was observed in the case of AMX+CLA. These observations highlight the highly efficient antimycobacterial activity of MEM.

Altogether, the susceptibility assays suggest that CwlM contributes to beta-lactam resistance, specifically to CTX and MEM. Nonetheless, we recognize that the observed changes in susceptibility may partially reflect impaired growth, rather than being solely attributable to differences in CW integrity.

The cytosolic NOD-like receptors (NLRs), NOD1 and NOD2, recognize molecules containing D-Glu-*m*-DAP and PG–derived muramyl dipeptide (MDP), respectively [7-8]. The intracellular survival assays showed that the depletion of CwlM resulted in a faster clearance of intracellular mycobacteria, possibly due to increased recognition of unmodified and uncross-linked PG muropeptides by THP-1-derived macrophages innate cytosolic receptors. Phosphorylated CwlM is known to stimulate PG biosynthesis by 30-fold [31], thus promoting the activity of mycobacterial PG-modifying enzymes such as NamH, MurT/GatD, and AsnB, as well as transpeptidases. It is inferable that *cwlM* repression impairs PG biosynthesis, leading to deficient PG modification and polymerization, thereby facilitating immune recognition by host cells. Since the depletion of CwlM was induced 24 h before infection, the induced *cwlM* KDM might have already been fragile at the time of infection. Although no considerable differences were observed in the OD_600_ values among the study strains, these could represent a population of compromised cells unable to withstand additional stress. Therefore, the induced *cwlM* KDM is likely more susceptible to intracellular stresses and macrophage-mediated antimicrobial killing, provoking a substantial survival drop shortly after internalization. This underscores the dual role of CwlM: not only is it essential for bacterial viability under basal conditions, but its depletion also curtails intracellular survival, thus highlighting its key role in sustaining survival within host macrophages. In conclusion, CwlM contributes to intracellular survival, perhaps by promoting efficient PG biosynthesis, and, in turn, enabling several mycobacterial PG-modifying immune evasion mechanisms.

The CwlM protein is annotated as a PG hydrolase that cleaves an amide bond between the glycan moiety (Mur*N*Ac) and the peptide moiety (L-alanine) [30]. However, several studies showed that CwlM lacks enzymatic activity, rather serving a regulatory function [31-32]. Here, we successfully cloned *cwlM* in frame with a C-terminal polyhistidine tag in pET29b and purified the His_6_-tagged CwlM_TB_ using AKTAprime. Afterwards, a zymogram was performed to determine whether CwlM_TB_ possesses amidase activity. PG from *M. luteus* is frequently used for the detection of amidase activity against the CW of gram-positive bacteria [30, 40]. In this case, the PG from *M. luteus* was used as a substrate because it enabled the assessment of PG-hydrolytic activity without the limitation provided by the external layers of the mAGP complex. Collectively, the zymogram and plate assays showed that CwlM_TB_ lacks PG-hydrolytic activity. Although our results contradict the findings of a previous study [30], they are concordant with the recent hypothesis proposing that CwlM plays a regulatory role in PG biosynthesis [31-32].

Building on the role of CwlM in PG biosynthesis, a recent study showed that the activity of PknB, the STPK for which CwlM is a major substrate, is stimulated by the depletion of D-*i*Glu amidation [14]. Accordingly, the depletion of D-*i*Glu amidation leads to perturbed PG cross-linking, hyperactivation of PknB and, ultimately, reduced *de novo* PG biosynthesis. The repression of *murT*/*gatD* results in the inhibition of Ldt activity and gives a crucial meaning to the activity of PBPs, which can be easily inhibited by beta-lactams [14]. Aligned with this, we have recently demonstrated that the depletion of MurT/GatD foments increased susceptibility to beta-lactams [6]. The sensing of uncross-linked PG by PknB promotes its overactivation and, consequently, stimulates the phosphorylation of its substrates, including that of CwlM, MurA, MurJ and others [14, 32, 61]. In this case, phosphorylated CwlM binds to FhaA, thereby stabilizing the activity of PbpA and inhibiting the activity of MurJ. Hence, lipid II is not translocated to the periplasm and accumulates in the cytoplasm, thereby reducing *de novo* PG biosynthesis [32]. Should the activity of PknB remain unregulated, this will eventually cause bacteriolysis.

Our work is not deprived of limitations. First, the CRISPRi-mediated repression of *cwlM* employed a medium-to-low strength PAM (PAM 14). Although the obtained levels of repression of *cwlM* were modest (about 3-fold), these were sufficient to cause observable viability defects. While our study aimed to investigate the role of *cwlM* in antibiotic susceptibility and intracellular survival, the observed growth impairment emphasizes the significance of CwlM for mycobacterial fitness. In fact, this near-lethal phenotype complicates the interpretation of drug susceptibility and intracellular survival data, as it may reflect the essentiality of *cwlM* for viability. Nonetheless, the observed loss of viability implies that *cwlM* is both essential and highly vulnerable, highlighting its potential as a therapeutic target. Future studies should thereby employ weaker PAMs for *cwlM* knockdown. Furthermore, the phosphorylation state of purified CwlM_TB_ remains unknown, as phosphorylation studies could not be conducted. Owing to its rapid growth, genetic homology, and conservation of core metabolic pathways, *M. smegmatis* (*Msm*) remains a widely used surrogate model for *Mtb*, particularly in exploratory studies such as this one [62, 63]. Indeed, *Msm* is a practical and insightful system for preliminary research on cell wall regulation and antibiotic susceptibility because of its higher permeability to beta-lactams, and its naturally occurring expression of conserved PG-modifying enzymes like CwlM. Nevertheless, we acknowledge that direct extrapolation of the obtained results to *Mtb* may be constrained due to differences in cell envelope composition, beta-lactamase activity, and virulence. Hence, additional assays are necessary to confirm the applicability of our findings in pathogenic *Mtb*.

## Conclusions

In conclusion, our findings suggest that CwlM may contribute to beta-lactam resistance and survival within host macrophages, thereby limiting antimicrobial efficacy and facilitating immune evasion. The essentiality of *cwlM* further underscores its potential as a promising therapeutic target. Taken together, the evidence presented here indicates that targeting CwlM-regulated processes could improve antibiotic sensitivity, shorten treatment regimens, and ultimately improve DR-TB patient outcomes.

## Declarations

### Availability of data and materials

The original contributions presented in the study are included in the main manuscript and in the provided supplementary material file, and further inquiries can be directed to the corresponding author and are available on reasonable request.

### Authors’ Contributions

C.S., D.M., F.O., E.A. and M.J.C. conceived and designed the study. C.S., D.M., and F.O. performed the experiments. C.S. created figure 3, D.M. and C.S. created and modified the rest, respectively. C.S., D.M., F.O., M.M., D.P., E.A. and M.J.C. analyzed the data. C.S. and D.M. wrote the original draft. All authors reviewed and approved the final version of the manuscript.

## Acknowledgments

The authors would like to thank the colleagues Tiago Gonçalves and Leonor Raposo at Filipa Vale’s Lab for providing us with the *Micrococcus luteus* strain and for sharing zymography protocols.

