## Supplementary Material for "Effects of CwlM, a peptidoglycan synthesis regulator, on beta-lactam resistance and host-pathogen interactions"

### 1 Supplementary Figures and Tables

#### 1.1 Supplementary Figures

AGATGAGATGGTCAGATTAACTGGCTGACGGAATTTATGCTTCTCCGACCATCAAGCATTTATCCGTACTCCTG
ATGATGCATGGTTACTCACCACTGCGATCCCCGGGAAAACAGCATTCCAGGTATTAGAAGAATATCCTGATTCA
GGTGAAAATATTGTTGATGCGCTGGCAGTGTTCCTGCGCCGGTTGCATTCGATTCCTGTTTGTAATTGTCCTTTT
AACAGCGATCGCGTATTTCGCCTCGCTCAGGCGCAATCACGAATGAATAACGGTTTGGTTGATGCGAGTGATTC
TGATGACGAGCGTAATGGCTGGCCTGTTGAACAAGTCTGGAAAGAAATGCATAATCTTTTGCCATTCTCACCGG
ATTCAGTCGTCACTCATGGTGATTTCTCACTTGATAACCTTATTTTTGACGAGGGGAAATTAATAGGTTGTATTG
ATGTTGGACGAGTCGGAATCGCAGACCGATACCAGGATCTTGCCATCCTATGGAACTGCCTCGGTGAGTTTTCT
CCTTCATTACAGAAACGGCTTTTTCAAAAATATGGTATTGATAATCCTGATATGAATAAATTGCAGTTTCATTTG
ATGCTCGATGAGTTTTTCTAATCAGAATTGGTTAATTGGTTGTAACACTGGCAGAGCATTACGCTGACTTGACG
GGACGGCGGCTTTGTTGAATAAATCGAACTTTTGCTGAGTTGAAGGATCAGATCACGCATCTTCCCGACAACGC
AGACCGTTCCGTGGCAAAGCAAAAGTTCAAAATCACCAACTGGTCCACCTACAACAAAGCTCTCACCAACCGTG
GCTCCCTCACGATATCAATAAACGAAAGGCTCAGTCGAAAGACTGGGCCTTTCGTTTATCTGTTGTTTGAAAAA
AAAAAGCGCCGCAACTGCGGCGCTTTTTTTTTTGAATTCTCTGACCAGGGAAAATAGCCCTCTGACCTGGGGAT
TTGCGATCCCTATCAGTGATAGATATAATCTGGGAACCCGCCGGTGACGCGTGAGCGTTTTTGTACTCGAAAGA
AGCTACAAAGATAAGGCTTCATGCCGAAATCAACACCCTGTCATTTTATGGCAGGGTGTTTTTTTTTTGTCGACT
TGGGGACCCTAGAGGTCCCCTTTTTTTTTTGA

**Figure S1 - Sequencing results (5' – 3') of PLJR962 with the sgRNA targeting the gene of interest (*cwlM*) in *M. smegmatis*.** The -10 site is underlined in yellow, the insert or sgRNA is underlined in green and the dCas9 handle is painted in grey.

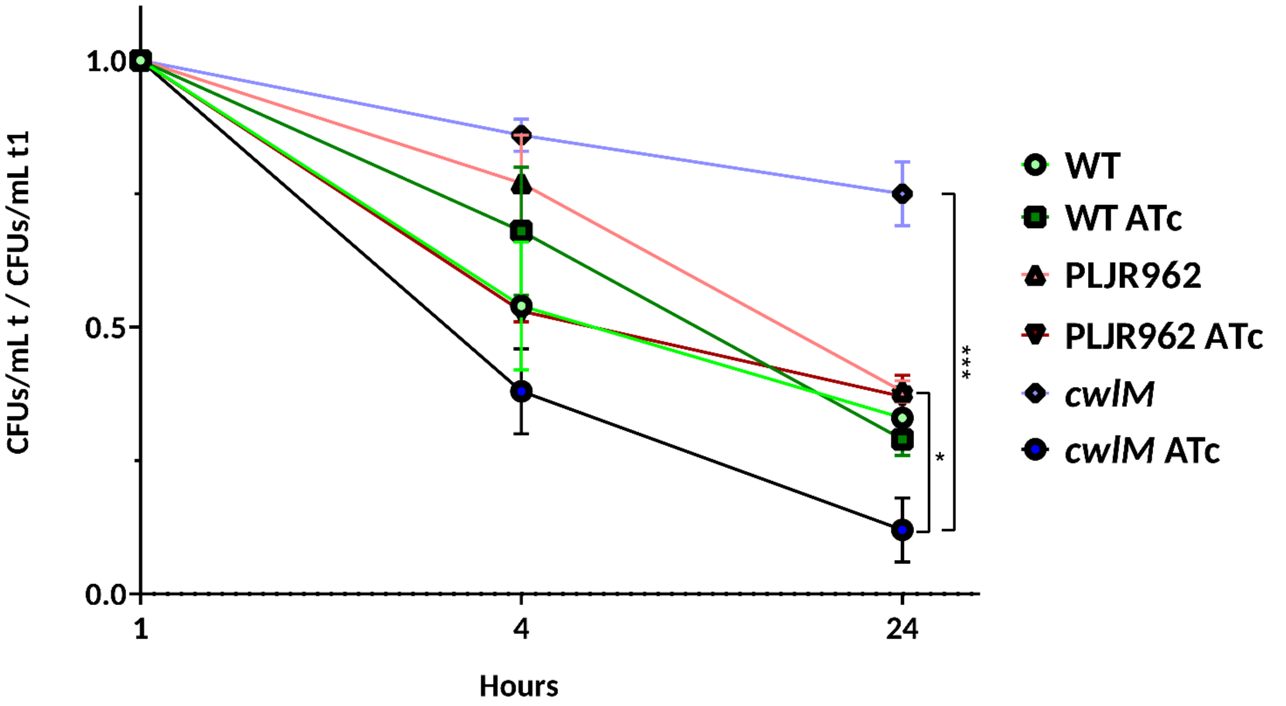


**Figure S2 – Ratio of bacterial survival (in CFUs/mL) in infected and disrupted PMA-differentiated THP-1 macrophages at 4 h and 24 h post-infection, compared to intracellular survival at 1 hour (t1) (n=3).** Comparative effect with and without 100 ng/mL of inducer (ATc). Error bars represent the standard error of the mean (SEM). Multiple comparisons were calculated using one-way ANOVA (* $P < 0.05$; *** $P < 0.001$).

**a. pET-29b:*cwlM* with primer Fwd T7:**

GGCGTACATTCCCTCTAGAATAATTTTGTTTAACTTTAAGAAGGAGATATACATATGCCGAGTCCGCGCCGCGAAGACG
GCGATGCGCTGCGCTGTGGCGACCGCAGTGCGGCCGTCACCGAGATCCGGGCTGCGCTGACCGCGTTAGGGATGCTG
GATCATCAGGAAGAAGACCTGACGACGGGCCGTAACGTCGCCCTTGAGTTGTTCGACGCGCAGCTCGACCAGGCGGTC
CGTGCCTTCCAACAGCATCGCGGCCTGCTGGTGGACGGCATCGTCGGTGAGGCCACCTACCGCGCGTTGAAAGAAGCC
TCCTACCGGCTCGGGGCCCGCACGCTGTACCACCAATTCGGCGCCCCGCTCTACGGGGACGACGTCGCTACACTGCAGG
CCCGGCTGCAGGATCTTGGTTTCTACACCGGGCTGGTCGACGGTCATTTCGGGTTGCAGACCCACAATGCGTTGATGTC
CTATCAGCGTGAGTACGGACTTGCCGCAGACGGTATCTGCGGCCCAGAAACGTTGCGCTCCTTGTACTTTCTAAGTTCG
CGAGTCAGCGGTGGCTCGCCACATGCGATTCGCGAAGAAGAGCTGGTCCGCAGCTCGGGGCCGAAGCTGTCTGGCAA
ACGGATCATCATTGATCCCGGTCGCGGCGGCGTGGACCACGGACTTATCGCGCAAGGTCCGGCTGGGCCCATCAGCGA
AGCAGACTTGTTGTGGGACTTGGCAAGTCGGCTCGAAGGACGGATGGCAGCTATCGGTATGGAGACCCACCTGTCCCG
TCCGACCAACCGTAGTCCGTCCGACGCAGAGCGTGCCGCCACCGCCAACGCCGTTGGCGCAGACCTGATGATCAGCCT
GCGCTGCGAGACCCAGACCAGTCTCGCGGCCAACGGCGTGGCTTCCTTTCACTTCGGCAACTCGCACGGCTCGGTGTCT
ACCATCGGCCGCAATCTTGCCGATTTCATTCAACGAGAAGTGGTGGCGCGCACCGGTTTACGGGATTGCCGTGTGCATG
GTCGAACGTGGGATCTGTTGCGGCTGACCAGGATGCCGACCGTTCAGGTCGATATCGGCTACATCACCAACCCCCACGA
TCGTGGGATGCTGGTCTCAACGCAGAACGC

**b. pET-29b:*cwlM* with primer Rv T7:**

AGCCTCGCTTCTTTCGGGCTTTGTTAGCAGCCGGATCTCAGTGGTGGTGGTGGTGGTGCTCGAGAGAACCGCCGAGTCT
ACCCGCTCGCTCGACAGACAGTTCGTGGGCCAGCAACTCGGCGAAAGTGAATGTGCCGGTGGGCCGATCGTTCTTGCCT
AACAGATACAGCCGTTTGACCGCGGCGAGAATGCCTTCGGCGATGGCATCGCGCGTCTGCGTTGAGACCAGCATCCCAC
GATCGTGGGGGTTGGTGATGTAGCCGATATCGACCTGAACGGTCGGCATCCTGGTCAGCCGCAACAGATCCCACGTTCG
ACCATGCACACGGCAATCCCGTAAACCGGTGCGCGCCACCACTTCTCGTTGAATGAAATCGGCAAGATTGCGGCCGATG
GTAGACACCGAGCCGTGCGAGTTGCCGAAGTGAAAGGAAGCCACGCCGTTGGCCGCGAGACTGGTCTGGGTCTCGCAG
CGCAGGCTGATCATCAGGTCTGCGCCAACGGCGTTGGCGGTGGCGGCACGCTCTGCGTCGGACGGACTACGGTTGGTC
GGACGGGACAGGTGGGTCTCCATACCGATAGCTGCCATCCGTCCTTCGAGCCGACTTGCCAAGTCCCACAACAAGTCTG
CTTCGCTGATGGGCCCAGCCGGACCTTGCGCGATAAGTCCGTGGTCCACGCCGCCGCGACCGGGATCAATGATGATCCG
TTTGCCAGACAGCTTCGGCCCCGAGCTGCGGACCAGCTCTTCTTCGCGAATCGCATGTGGCGAGCCACCGCTGACTCGC
GAACTTAGAAAGTACAAGGAGCGCAACGTTTCTGGGCCGCAGATACCGTCTGCGGCAAGTCCGTACTCACGCTGATAGG
ACATCAACGCATTGTGGGTCTGCAACCCGAAATGACCGTCGACCAGCCCGGTGTAGAAACCAAGATCCTGCAGCCGGGC
CTGCAGTGTAGCGACGTCGTCCCCGTAGAGCGGGGCGCCGAATTGGTGGTACAGCGTGCGGGCCCCGAGCCGGTAGG
AGGCTTCTTTCAACGCGCGGTAGGTGGCCTCACCGACGATGCCGTCCACCAGCAGGCCGCGATGCTGTTGGAAGGCACG
GACCGCCTGGTCAAGCTGCGCGTCAAAAAC

**Figure S3 - Sequencing results (5' – 3') of pET29b with the T7 promoter a) forward and terminator b) reverse primers**. Fwd – forward, Rv – reverse. The insert is underlined in green; the histidine tail is underlined in purple; and the XhoI and NdeI restriction sites are underlined in grey and light blue, respectively.

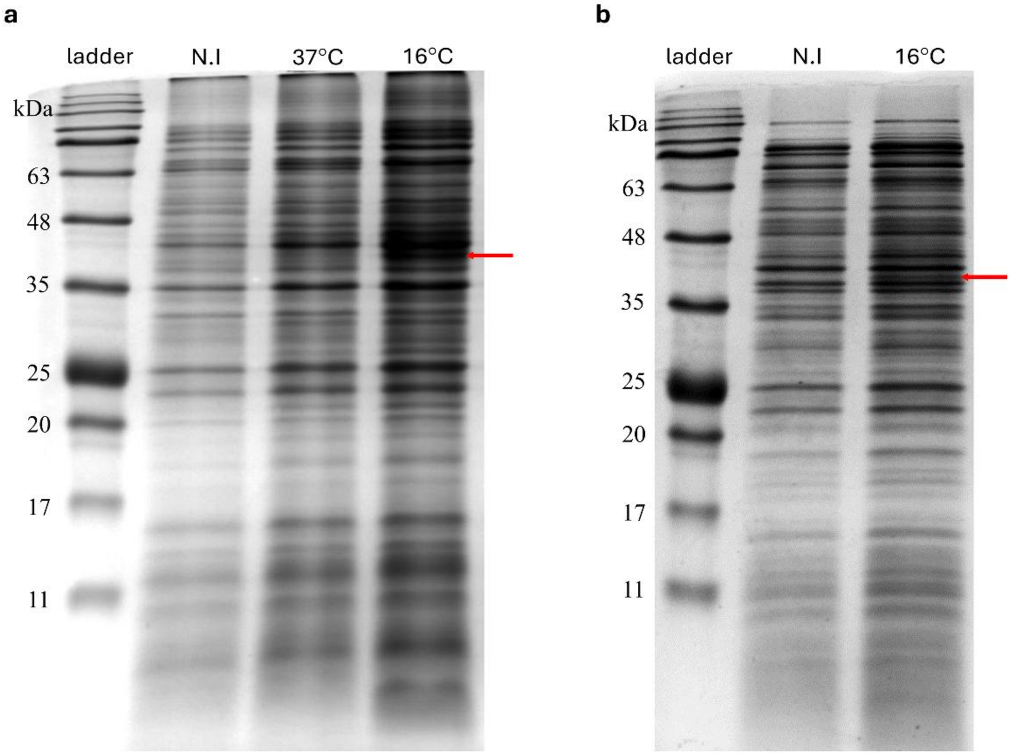


**Figure S4 - SDS polyacrylamide gel analysis of cell lysate supernatants for the optimization of the expression and induction conditions for the purification of the CwlM protein.** Protein separation is based on size and 10% acrylamide gels were used. The size of CwlM is predicted to be about 43.9 kDa. The red arrows indicate the presence of the protein of interest. **(a)** SDS-PAGE of the supernatants of non-induced and induced cell lysates of *E. coli* BL21 (DE3) expressing CwlM in different conditions. Lane order: NZYColour Protein Marker II ladder; non-induced (N.I) *E. coli* BL21(DE3):pET-29b:*cwlM*; *E. coli* BL21(DE3):pET-29b:*cwlM* induced at 37°C for 3 h; *E. coli* BL21(DE3):pET-29b:*cwlM* induced at 16°C overnight. **(b)** SDS-PAGE of the supernatant of the non-induced and induced cell lysates of *E. coli* BL21 (DE3) expressing CwlM at 16°C, overnight**.** Lane order: ladder NZYColour Protein Marker II ladder; non-induced (N.I) *E. coli* BL21(DE3):pET-29b:*cwlM*; and *E. coli* BL21(DE3):pET-29b:*cwlM* induced at 16°C overnight.

### 1.2 Supplementary Tables

**Table S1 - Average of three independent experiments measuring the optical density** (at 600 nm) **of the study strains** (*M. smegmatis* WT, PLJR962 and the *cwlM* knockdown mutant, grown with and without 100 ng/mL of ATc) **immediately before infection.**

| **Optical density (600 nm) immediately before infection** | |
|---|---|
| *M. smegmatis* | |
| WT | 0.57 |
| WT ATc | 0.50 |
| PLJR962 | 0.41 |
| PLJR962 ATc | 0.43 |
| *cwlM* | 0.34 |
| *cwlM* ATc | 0.41 |
